# Anterior thalamic nuclei: A critical substrate for non-spatial paired-associate memory in rats

**DOI:** 10.1101/2021.08.14.456360

**Authors:** Jennifer J Hamilton, John C Dalrymple-Alford

**Affiliations:** School of Psychology, Speech and Hearing, University of Canterbury, 20 Kirkwood Avenue, Christchurch 8041, New Zealand; New Zealand Brain Research Institute, 66 Stewart Street, Christchurch 8011, New Zealand; Brain Research New Zealand – Rangahau Roro Aotearoa

**Keywords:** thalamic amnesia, biconditional discrimination, extended memory system, rat, immediate early genes.

## Abstract

Injury or dysfunction in the anterior thalamic nuclei (ATN) may be the key contributory factor in many instances of diencephalic amnesia. Experimental ATN lesions impair spatial memory and temporal discriminations, but there is only limited support for a more general role in non-spatial memory. To extend evidence on the effects of ATN lesions, we examined the acquisition of bi-conditional associations between odour and object pairings presented in a runway, either with or without a temporal gap between these items. Intact adult male rats acquired both the no-trace and 10-second trace versions of this non-spatial task. Intact rats trained in the trace version showed elevated Zif268 activation in the dorsal CA1 of the hippocampus, suggesting that the temporal component recruited additional neural processing. ATN lesions completely blocked acquisition on both versions of this association-memory task. This deficit was not due to poor inhibition to non-rewarded cues or impaired sensory processing, because rats with ATN lesions were unimpaired in the acquisition of simple odour discriminations and simple object discriminations using similar task demands in the same apparatus. This evidence challenges the view that impairments in arbitrary paired-associate learning after ATN lesions require the use of multimodal spatial stimuli. It suggests that diencephalic amnesia associated with the ATN stems from degraded attention to stimulus-stimulus associations and their representation across a distributed memory system.

## Introduction

The anterior thalamic nuclei (ATN) have complex reciprocal connections with many brain structures in a ‘hippocampal-diencephalic-cingulate’ network (Bubb et al., 2017). The pattern of connections suggests that this small brain region is a critical subcortical node that supports memory. This perspective aligns with evidence that loss of ATN neurons in people with the Korsakoff’s syndrome, which is characterised by severe and persistent amnesia, discriminates these patients from alcoholic individuals who retain relatively good memory (Harding et al., 2000; Kopelman, 2015; Maillard et al., 2021). Intra-thalamic recordings in epilepsy patients also reinforce the association between ATN function and memory (Sweeney-Reed et al., 2021). Additional support comes from disruption to the ATN caused by a mammillothalamic tract (MTT) disconnection (Dillingham et al., 2019; Perry et al., 2018), which is the brain injury that most consistently corresponds with memory impairment after thalamic infarcts (Carlesimo et al., 2011).

The association between the ATN and memory is supported by a large body of experimental lesion studies. These studies, however, have overwhelmingly focused on spatial memory and the similarity between the effects of ATN and hippocampal lesions (Aggleton & Brown, 1999; Aggleton & Nelson, 2015; Dalrymple-Alford et al., 2015; Nelson, 2021; Perry et al., 2018). However, the clinical evidence shows that memory dysfunction extends beyond spatial information (Rempel-Clower et al., 1996; Turriziani et al., 2004). The existence of non-spatial deficits after experimental ATN lesions has some support. ATN dysfunction slows rats’ ability to form an attentional set when learning a series of different non-spatial stimuli pairings in which only one stimulus dimension is rewarded (Wright et al., 2015; Bubb et al., 2021; Nelson, 2021). Latent inhibition associated with prior exposure to non-reinforced auditory cues is also impaired (Nelson et al., 2018). Conversely, relational processing of nonspatial stimuli during sensory preconditioning is unimpaired by ATN lesions (Ward-Robinson et al., 2002). Evidence from tasks that focus more explicitly on memory has thus far been limited to recall of the temporal order of non-spatial items, but these effects are limited by specific task demands. That is, ATN lesions disrupted relative recency judgements when a list of multiple odours or multiple objects was presented in a single block of trials, but not when relative choice was based on non-spatial stimuli presented across more distinct temporal episodes, including a single pair of items or two blocks of multiple items (Wolff et al., 2006; Aggleton et al., 2011; Dumont & Aggleton, 2013; Mitchell & Dalrymple-Alford, 2005). Clearly, more examples of ATN-lesion deficits in non-spatial memory are needed to support the specific association between clinical amnesia and memory impairment after ATN dysfunction.

Failure to acquire memory for an association of arbitrary stimuli is a core feature of the amnesic syndrome (Turriziani et al., 2004). These tasks offer an opportunity to explore non-spatial memory. One class of association memory is the ability to discriminate a combination of cues relative to their component elements (McDonald et al., 1997). Like hippocampal lesions, ATN lesions do not impact these configural learning tasks when they rely on non-spatial cues (Chudasama et al., 2001; McDonald et al., 1997; Moran & Dalrymple-Alford, 2003; Ridley et al., 2002). Conditional learning tasks provide a second class of arbitrary associations. This time, one correct response is signalled by one stimulus and a second correct response is signalled by an alternate stimulus. Unlike hippocampal lesions, however, ATN lesions did not delay acquisition in these tasks when either egocentric responses or non-egocentric spatial choices were signalled by a single visual cue (Sziklas & Petrides, 2004, 2007).

Paired-associate learning tasks offer an alternative test of memory for arbitrary associations. These memory tasks require the formation of unique stimulus-stimulus representations. This is different to the situation when a discrimination is based on a combination of cues contrasted with their individual elements or when the conditional relationship is determined by memory a single salient cue and its consequence. Paired-associate tasks often use stimulus pairings that require a biconditional relationship among two stimuli that have separate attributes (different types of information). For example, an odour attribute combined with a location attribute is rewarded (e.g. odour 1 + location A; and odour 2 + location B), but the alternate pairings are not rewarded (odour 1 + location B; and odour 2 + location A). Like hippocampal lesions (Gilbert & Kesner, 2002, 2003; Hunsaker et al., 2006; Jo & Lee, 2010; Lee & Solivan, 2008; Sziklas & Petrides, 2002), ATN lesions produce severe acquisition deficits when one of the attributes concerns location, such as in odour-place or object-place association tasks (Dumont et al., 2014; Gibb et al., 2006; Sziklas & Petrides, 1999). These impairments are not solely due to the inclusion of a place attribute, because ATN lesions had far milder effects on non-conditional spatial discriminations using similar test procedures (Dumont et al., 2014; Gibb et al., 2006). However, ATN lesions did not impair biconditional discriminations when local contextual cues were used that were either visual, thermal or texture (Dumont et al., 2014). These findings suggest that the inclusion of spatial attributes may be necessary to reveal impairments in learning arbitrary associations after ATN lesions.

We have, however, found preliminary evidence of impaired acquisition of a non-spatial object-odour association task after ATN lesions (Bell, 2007). This study used an open circular cheeseboard platform (Gilbert & Kesner, 2003), so it is possible that task characteristics introduced spatial elements that disrupted acquisition. The current study, therefore, examined acquisition of non-spatial associative memory in a high-walled red-Perspex runway that minimised the availability of spatial cues. The runway also enabled us to assess the influence of an explicit 10-second temporal delay (i.e. a trace) between the presentation of the odour and object stimuli. Kesner and colleagues have suggested that hippocampal lesions do not prevent the acquisition of object-odour associative memory unless the procedure includes an explicit temporal gap between the nonspatial stimuli, which was linked with the specific loss of dorsal CA1 neurons (Gilbert & Kesner, 2002; Kesner et al., 2005). Both ATN lesions and mammillothalamic tract lesions diminish the number of CA1 dendritic spines (Harland et al., 2014; Dillingham et al., 2019) and can reduce hippocampal immediate early gene (IEG) activation (Jenkins et al., 2002; Jenkins et al., 2002; Dupire et al., 2013; Loukavenko et al., 2015; Perry et al., 2018). We anticipated that ATN lesions would slow the acquisition of the standard odour-object paired-associate learning task in the runway when there is no explicit trace between the paired stimuli, but severely impair acquisition in the trace version of this task because of the additional temporal component. We complemented this work by examining Zif268 IEG expression in memory-related brain structures after a retention session conducted 5 days after the end of training.

## Materials and Methods

### Animals and housing conditions

Male Long-Evans rats (12 months old; average 630g) were housed in groups of three of four in Makrolon cages (48 x 28 x 22 cm) on a reversed 12-hour light-dark cycle (lights on at 8pm). Behavioural testing occurred in the dark phase of the light cycle, five sessions per week; observations confirmed that rats were relatively more active during the dark phase. Rats were maintained at 85% of free feeding body weight with water ad-libitum; food was ad-libitum prior to and after surgery for post-operative recovery. All procedures complied with approved guidelines from the University of Canterbury Animal Ethics Committee. Four groups of rats were established by random assignment after pre-surgery familiarization in the radial arm maze (RAM) and runway. The four groups were bilateral ATN lesions with or without a trace between odour and object stimuli when trained in the paired associate task and the corresponding two Sham-lesion groups. The final sample sizes were: (1) Sham-No Trace = 7 (that is, a Sham-lesion group with no interval between odour and object stimuli used in the paired-associate task); (2) Sham-Trace = 7 (Sham-lesion group given a 10 second trace between presentation of the odour and the object stimuli); (3) ATN-No Trace = 8; and (4) ATN-Trace = 9. An additional ATN-lesion rat was excluded for not meeting the lesion criteria (see below).

### Surgery

Rats were anaesthetised with isoflurane (induction 4%; maintenance ∼2.5%) and placed in a stereotaxic apparatus with atraumatic ear bars (Kopf, Tujunga, CA). The incisor bar was set to −7.5mm below interaural line to minimise fornix damage. In each hemisphere, two infusions were directed at the anteroventral nucleus (AV; upper and lower) and one infusion was directed at the anteromedial nucleus (AM). Each surgery used one of five anterior-posterior (AP) coordinates relative to an individual rat’s Bregma-Lambda distance (all values in mm). For AV lesions, the AP coordinates were: −2.20 for B–L 6.9–7.2; −2.25 for B–L 7.3 to 7.6; −2.30 for B-L ≥ 7.7. The AV infusions were made at −5.68 and −5.73 below dura and ±1.52 lateral to the midline. The AM infusion was made 0.1 mm more anterior than the AV coordinate, at −5.76 below dura and ±1.16 lateral; the first AM lesion was made after completing all AV infusions and began contralateral to the last AV lesion site. 0.15M N-methyl-d-aspartate (NMDA; Sigma, Castle Hill, NSW) in 0.1M phosphate buffer (pH 7.20) was infused into each site at 0.04ul per minute via a 2.5μL Hamilton syringe (Reno, NV, USA) using a micro infusion pump (Stoelting, Wooddale, IL). The AV sites per hemisphere received 0.12 and 0.10μL for the dorsal and ventral sites, respectively; 0.06μL was used for each AM site. The syringe needle remained in situ for 3 minutes per site, post-infusion. Sham lesion surgeries used the same procedure but the needle was lowered to 1 mm above the upper AV and the AM lesion sites with no infusion given.

### Behavioural Tasks

An 8-arm RAM was used to confirm the spatial memory impairment in ATN-lesion rats following surgery. A runway apparatus was used to train the rats on all non-spatial tasks; i.e. simple odour and simple object discrimination, and odour-object and odour-trace-object paired-associate tasks. Prior to surgery, food-restricted rats were habituated to the 8-arm RAM, described below, to retrieve 0.1g chocolate pieces (FoodFirst LTD Auckland, NZ) and adapt to the opening and closing of arm doors. In a red Perspex runway (Fig. 1), rats were also shaped to nose poke a 2cm circular white plastic cap that was 10cm above floor level and at the centre of a thin sponge on a clear Perspex insert that was attached to the end wall. They were also shaped to push (using nose or paw) an object hinged at the rear of its base to the top of a food well that was recessed at the centre of a 5cm x 8.5cm x 3cm wooden block. The food well contained chocolate pieces (and held inaccessible food beneath the recess). The object used for pretraining was not used again. Following surgery recovery rats were re-familiarised to both the RAM and the runway.

**Figure 1.**
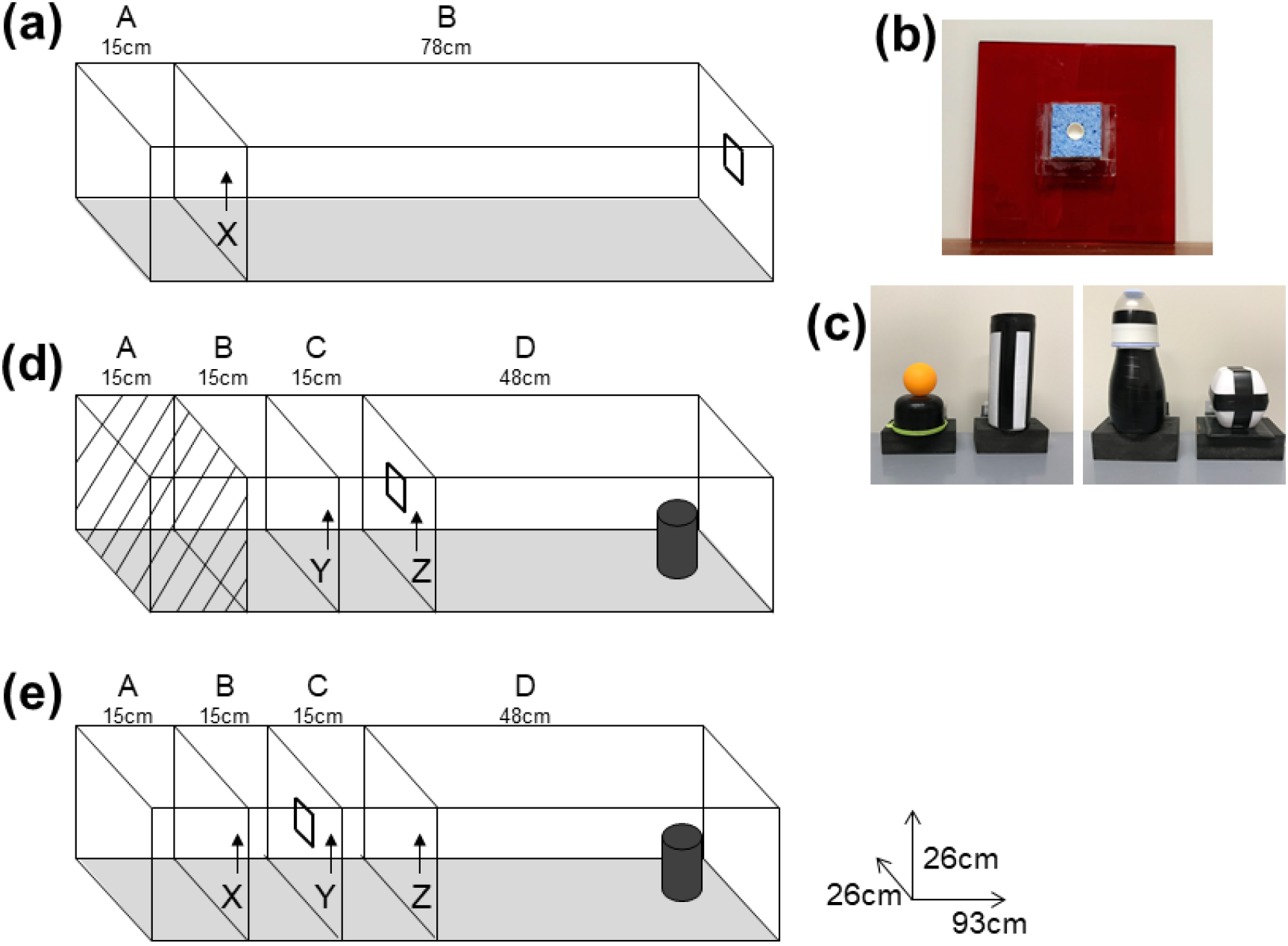
Runway apparatus used in both the simple odour and simple object discrimination tasks, and the odour-object paired associate memory tasks. **(a)** For simple discriminations, rats were placed in the start box (A), door X was removed and we recorded the latency to search for food in the receptacle inside the odourised sponge **(b)** or search for food by pushing back a single object located at the end of compartment B **(c)**. Runway configuration for No Trace (**d**) and Trace (**e**) paired-associate memory tasks. **(d)** Rats began the No Trace paired-associate task in Compartment B for 120 seconds on the first trial on any session and for 20 sec on the remaining trials (Compartment A was not used). Door Y was removed to give access to the odour on door Z at the end of Compartment C, per **(b)**. Immediately after the food reward was consumed from the plastic cap in the centre of the odour sponge, Door Z was removed and we recorded the latency to interact with a single object, **(c),** at the end of Compartment D. **(e)** Rats began the Trace paired-associate memory task in Compartment A for 120 seconds on the first trial per day and 20 secs on remaining trials. Door X was removed to give access to the odour sponge, per **(b)**, on door Y at the end of Compartment B. Door Y was immediately removed on eating the food and the rat was retained for a 10-second trace (delay) in Compartment C before Door Z was removed. For the paired-associate tasks, we recorded latency from removal of Door Z to interaction with the object at the end of Compartment D.

### Spatial working memory in the radial arm maze

Post-surgery, spatial working memory was tested using the RAM located in the centre of a windowless room (3 x 3m). The maze had a grey-painted wooden 35cm wide central hub with 8 aluminium arms (65cm long x 8.6cm wide, with 4.5cm-high borders), raised 67.5cm above the room floor. Clear Perspex walls (19cm long x 25cm high) extended along one side of each arm from the hub to prevent rats from jumping between arms. A black wooden food well (5cm x 8.5cm x 3cm), with inaccessible food underneath, was placed at the end of each arm. During testing, 2 x 0.1g chocolate pieces were present in each food well at the start of each trial. Clear Perspex guillotine doors, which could be raised up from beneath the hub by a pulley system, controlled access to the hub and arms.

Ten consecutive days of formal training in the RAM was used to test spatial working memory. The rat was placed in the central hub and ∼10 seconds later all eight arms were opened and the rat allowed to make a choice defined as both hind feet over an arm threshold. Once the rat had entered an arm, all the doors were closed and it was confined to the arm for ∼15 seconds irrespective of whether it made a correct choice. The rat was then allowed back into the central hub and held there for ∼10 seconds before all the doors were re-opened to allow another arm choice. The trial concluded when the rat had visited all 8 arms, 20 arm choices had been made or 10 minutes had elapsed.

### Non-spatial tasks: Simple discrimination and paired-associate learning

Both simple discrimination and paired-associate tasks used the same red Perspex runway (93cm x 26cm x 26 cm) placed on a 75cm high table (Fig. 1). The runway could be divided by three vertically removable doors (red Perspex, 26cm x 26cm) that were selected on the basis of the specific behavioural task. As for familiarisation, the 6.5cm x 6cm x 8mm thin sponge with the cap at its centre held a chocolate food reward (Fig. 1). The sponge was attached to a removable door in the runway (for the paired-associate tasks) or to the end wall of the runway (for the simple discrimination task). A different door plus sponge was used for each odour, which was infused adjacent to the receptacle with 5ml of sunflower oil mixed with 20µl of an Essential Oils of New Zealand substance. The odours were lemon and clove for the paired associate tasks, and cinnamon and lime for the simple discrimination tasks, or the reverse in counterbalanced groups. Light and visually distinct objects (Fig. 1) were attached by a hinge at the rear of its base to the top of the food well, so that a rat could easily push the object back (either nose or paw) to search the well for food reward. A wooden frame (1.8m high from floor x 1.3m wide), draped in black curtains, enclosed all sides of the test area to reduce spatial cues during testing.

The simple discriminations (odour only; object only) and the paired-associate tasks used a ‘go/no-go’ procedure. The first trial for each session began by restraining the rat in the start area for 120 seconds to reacclimatise to the maze; subsequent trials used 20 seconds in this start area. All rats received 12 massed trials per daily session, with 6 go (rewarded) and 6 no-go (non-rewarded) trials in a pseudo-randomised order. No more than three go or three no-go trials were run consecutively in a session. None of the four different pairs of odour-object stimuli was repeated on consecutive trials. Correct items in the simple discrimination tasks and correct stimulus pairs in the paired-associate tasks were counterbalanced across rats. For the simple odour discrimination, a response was defined as a nose poke into the cap in the centre of the odourised sponge; sniffing the sponge only was considered a no-go response. Food was always present in the cap at the centre of the sponge for the odour-object paired-associate tasks; the go / no-go aspect here refers to whether the object was pushed to reveal whether a food reward was under it, at the end of the runway. For objects, a response was defined as any push with nose or paw that tilted the object backwards. Latency from the time that the last door was lifted and when the rat interacted with the odour (nose poke in the cap; for simple odour discrimination) or with the object (pushed back on hinge, for simple object discrimination and paired-associate tasks) was measured using a quiet stopwatch (Dick Smith, New Zealand). The observer was seated within the curtains and adjacent to the runway to operate the doors and record behaviour. A trial was designated as “correct” if the rat responded in less than 8 seconds on go trials or did not respond before 8 seconds on no-go trials.

### Simple discriminations

Half of the rats received training in the simple odour discrimination task followed by the simple object discrimination and vice versa for the other half. Latency between door X (Fig. 1a) and interaction with the odour or object at the end of the runway was recorded. Rats were trained on the simple odour and simple object discrimination task until they reached criterion (80% correct over 2 consecutive days). For the simple odour discrimination task, a single odour was presented on each trial. The rat had to discriminate which of two odours was paired with a food reward presented in the sponge at the end of Compartment B (Fig. 1a). In the simple object discrimination task, a single object was presented on each trial at the end of Compartment B (not shown in Fig. 1). The rat had to learn which of two objects was paired with food reward presented under the object.

### Paired-associate tasks (trace and no-trace)

These tasks used a new layout of vertically removable doors (Fig. 1). The rats learned which of two odour-object pairings would be rewarded (e.g. odour 1 + object A; and odour 2 + object B) and which of two pairings were not rewarded (i.e. odour 1 + object B; and odour 2 + object A). Irrespective of the pairing, rats always received a reward on presentation of the odour when allowed into the second compartment of the runway. This was compartment C for the “no-trace” condition in which compartment A was not used and compartment B was the start compartment. The odour was at the end of compartment B for the “trace” condition, for which compartment A was used as the start box. Rats in the trace condition were subjected to a 10 second delay between presentations of the odour and object stimuli, by being held in compartment C, whereas rats in the no-trace condition were exposed to the object immediately following their interaction with odour (after eating the reward). In both trace and no-trace conditions, we recorded latency to traverse compartment D, from the time door Z was lifted until the rat interacted with the object or refrained from interacting with it for 8 seconds. Two groups of rats (Sham-lesion and ATN-lesion) were trained on the odour-object paired-associate tasks (No Trace groups) and two corresponding groups were trained on the odour-trace-object paired-associate task (Trace groups) until they reached criterion (80% correct trials across 3 days) or a maximum of 50 days.

### Post-acquisition retention test

Retention of the paired-associate task for any rat was conducted 5 days after reaching criterion or after 50 days of training. This session used 12 massed rewarded/non-rewarded trials on a single day in identical fashion as used previously for that rat.

### Histology

#### Perfusion and tissue collection

On the 3 days prior to the retention test, single rats were familiarised to an empty clean cage for 90 minutes in a dark quiet room. Immediately following the last trial in the retention test, the rat was placed in its cage in the familiar dark quiet room 90 minutes prior to perfusion. Rats were deeply anaesthetised with sodium pentobarbital (125 mg/kg) and perfused transcardially with saline followed by paraformaldehyde (4% PFA in 0.1M phosphate buffer; PB) to fix the brain. Brains were post-fixed in 4% PFA followed by a minimum of 48 hours in a long-term solution (20% glycerol in 0.1M PB). Coronal 40µm sections were collected using a freezing microtome (Thermofisher, UK) and stored in a cryo-protectant solution (30% glycerol, 30% ethylene glycol in 0.1M PB) at −20°C until processed for immunohistochemistry. The coronal sections were collected in two separate series. The first series captured consecutive sections in 5 cryovials for immunohistochemistry (i.e. 1 in 5). This series included an anterior block of sections from the prefrontal cortex to the septal area (approximately +4.7 to −0.6mm from Bregma) and a posterior block from the mediodorsal thalamus (approximately −3.5mm from Bregma) to the caudal retrosplenial cortex. The second series captured consecutive sections for lesion verification. These sections were collected in 4 cryovials from immediately anterior to the ATN to the posterior mediodorsal thalamus (approximately −0.6 to −3.5mm from Bregma).

#### Neu-N staining and ATN lesion verification

Free floating sections throughout the ATN region were washed (3 × 10 min) in 0.1M phosphate buffered saline with Triton-X (0.2%; PBSTx) before incubation in endogenous peroxidase blocking buffer for 30 minutes (1% hydrogen peroxide (H_2_O_2_), 50% methanol (CH_3_OH) in 2% PBSTx). Sections were incubated overnight at 4°C in anti-NeuN primary antibody (1:5000; monoclonal-Mouse Cat# MAB377; Millipore, California USA) in PBSTx with 1% normal goat serum (NGS; Life Technologies, NZ). Excess antibody buffer was removed with PBSTx and followed by incubation in biotinylated goat anti-mouse secondary antibody (1:1000 Cat# BP-9200-50: Vector Laboratories, California USA) overnight at 4°C in PBSTx and 1% NGS. Sections were placed in ExtrAvidin (peroxidase conjugated; 1:1000; Sigma, NSW Australia), PBSTx and 1% NGS for 2 hours at room temperature. To remove excess ExtrAvidin and Triton X-100, the sections were washed in PBS, PB, and then Tris buffer (pH 7.4 in distilled H_2_0), to prepare the sections for visualisation with freshly prepared diaminobenzidine (DAB 0.05%; Sigma, in 0.01% H_2_O_2_ in Tris buffer). The DAB reaction (approximately 5 min) was stopped using Tris buffer (1 × 10 min wash) and sections were placed in PB at 4°C overnight before mounting on gelatinised slides and allowed to dry. The slides were dehydrated through graded alcohol (70-100%) before being cleared in xylene and mounted with DPX (06522; Sigma Aldrich) and a coverslip.

The number of Neu-N positive cells in the ATN was used to determine lesion extent. This used sections from both the left and right hemisphere for every one in four 40µm sections throughout the ATN and the area quantified using ImageJ (NIH, USA). NeuN-positive cell staining was photographed at 5x objective on a Leica DM6 B upright microscope and DFC7000T camera (Leica Microsystems, Germany). Automated counts of the cells were obtained through ImageJ (image analysis software, National Institute of Health, NIH, USA). The area surrounding neurons in the ATN region was manually selected and the images were converted to 8-bit grey scale, background was subtracted (rolling = 40), converted to mask and the watershed function was applied, and all neuronal cells above threshold (‘MaxEntropy’ threshold, circularity 0.5-1.0) were counted. The detection threshold was the same for all sections. Automated NeuN-stained nuclei of cells were counted in nine of the sham rats (5 in the Sham-Trace group and 4 in the Sham-No Trace group) to express cell counts in the ATN. Automated counts of NeuN-stained nuclei of spared cells in all ATN lesion rats were compared relative to the average count of NeuN cells observed in the sham rats. Acceptable lesions were defined as reaching the criterion of 50% loss of cells in the ATN, with a minimum of 25% per hemisphere. This is consistent with ATN lesion criteria in previous studies (Mitchell & Dalrymple-Alford, 2005, 2006; Perry, et al., 2018).

#### Zif268 immunohistochemistry

Free floating sections were processed in a similar way as for NeuN, but incubated in rabbit polyclonal Zif268 primary antibody (Egr-1; 1:1000 Cat# sc-110; Santa Cruz Biotechnology, USA) for 72 hours at 4°C in PBSTx with 1% NGS, followed by incubation in biotinylated goat anti-rabbit secondary antibody (1:1000: Vector Laboratories BA-1000) overnight in PBSTx and 1% NGS. Following DAB visualisation (∼18 minutes), Zif268 positive cell staining in each region of interest was photographed with a 10x objective with a light microscope (Leica, Germany). Automated counts of the cells were obtained through ImageJ (NIH, USA) in the same manner as NeuN-positive cells, except circularity was set at 0.65-1.0. Counts of Zif268-positive cells for all regions used the same threshold algorithm, with between two to six sections per region of interest in each rat quantified. The average Zif268 positive cell count per mm^2^ across sections (from both hemispheres) within a region of interest was used.

#### Data analysis

ANOVA using Statistica (v13; Dell Inc.) was conducted to test mean differences across the four groups of rats (Sham-No Trace; Sham-Trace; ATN-No Trace; and ATN-Trace). Repeated measures factors were added for Days (RAM and simple discrimination), Blocks of Trials (paired-associate tasks), and Zif268 counts across related regions or subregions of interest. The ANOVA for both the RAM and simple discrimination tasks compared four groups to confirm that the two Sham-lesion and two ATN-lesion groups were consistent with each other, but note that there was no trace component in the RAM or simple discrimination tasks. For both simple discrimination and paired-associate tasks in the runway, a reciprocal transformation of latency data in individual trials was used to ensure homogeneity of variance. On any given test day, the transformed latencies generated a mean latency for each of the six non-rewarded trials and the six rewarded trials. For the simple discrimination tasks, we analysed the mean latency difference score (the average of rewarded trials minus the average of the non-rewarded trials) but not the rewarded and non-rewarded trials separately due to a data storage failure. All three latency measures were available for the paired associate memory tasks (non-rewarded trials; rewarded trials; and latency difference). To account for multiple comparisons and to balance Type I and Type II errors we used the significance level of p<0.02 for behavioural analyses as there were three related measures in the paired-associate task (non-rewarded trials, rewarded trials, and mean latency difference). There were 6 primary analyses for regions of interest for Zif268 expression, so significance in these analyses was set at p<0.01. Post-hoc Newman-Keuls (N-K) tests assessed pairwise group differences. Simple main effects analysis was used when there were significant interactions involving repeated measures factors.

## Results

### Lesion verification

All but one of the 18 rats with ATN lesions met the a priori criterion of 50% bilateral damage to the ATN and a minimum of 25% per hemisphere. The median bilateral damage was 76% across both ATN-lesion groups (range: ATN-No Trace, 61-93%; ATN-Trace, 63-93%; Fig. 2). The excluded rat had a satisfactory lesion in one hemisphere only (48% total, 75% left, and 20% right). Figure 2d shows that the NeuN counts for ATN neurons in the ATN-lesion rats (M = 3388, SD = 2024) were all well below the values found in the sham-lesion rats (M = 15339, SD = 1065; this count was similar to that reported by Frost et al. 2020). Eight of the 17 lesion rats (4 in the Trace group and 4 in the No-Trace group) had no damage to either the mediodorsal thalamus (MD) or reuniens/rhomboid (Re/Rh) nuclei and there was relatively little damage to these structures in the remaining 9 rats with lesions. When MD damage occurred, it was in the anterior aspects that lie adjacent to the ATN. So, MD damage was assessed from approximately +1.72 to +2.10 mm from Bregma, where it ranged from 0-41% loss of cells across the whole group (median = 21%). When Re/Rh damage occurred it was also adjacent to the ATN region, so it was assessed approximately −1.08 to-2.04 mm from Bregma, where it ranged from 0-33% (median=6%). Note, these percent injury values do not represent the entire MD and Re/Rh regions, because the posterior regions of both nuclei were intact in all ATN-lesion rats.

**Figure 2.**
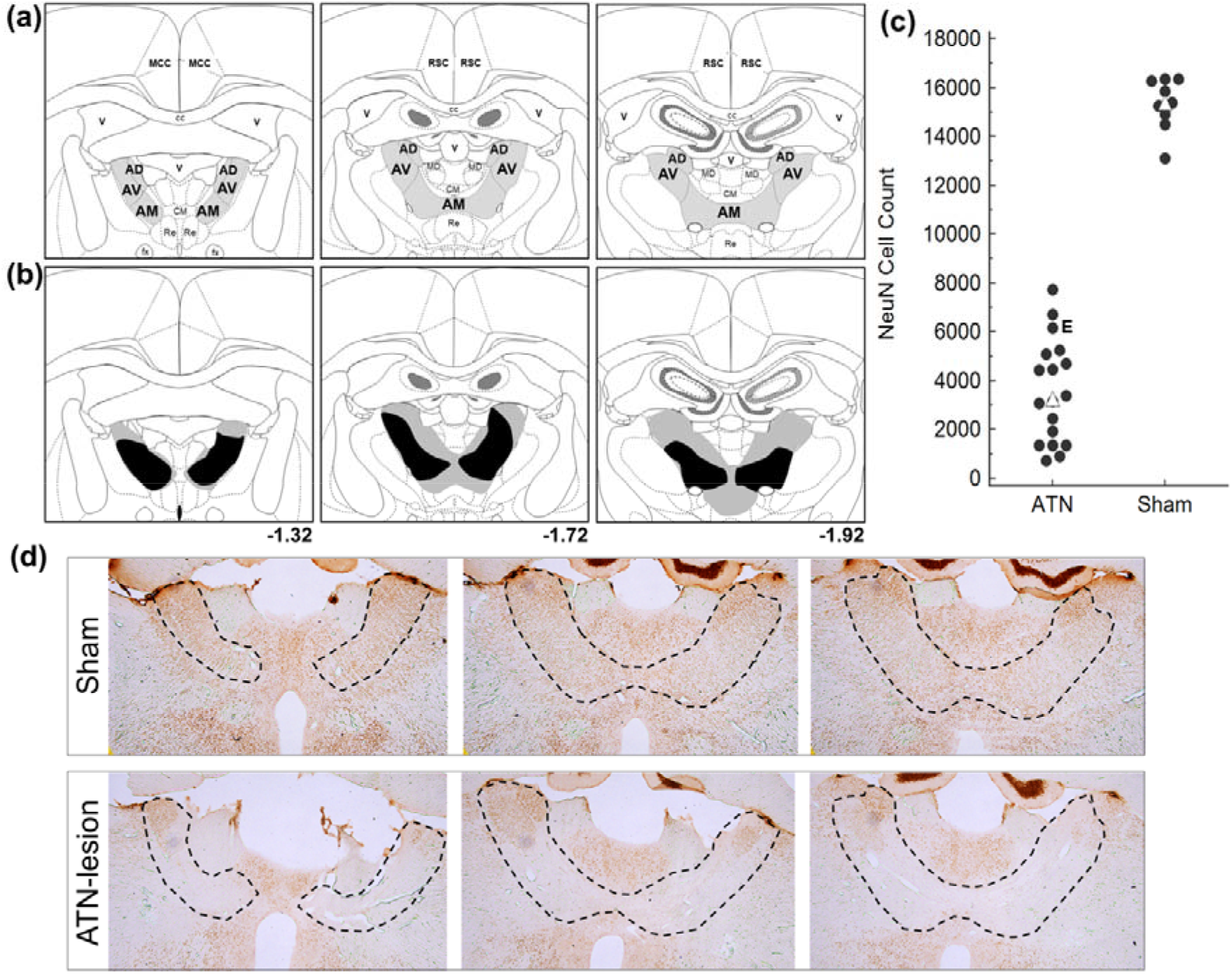
Schematic figure of **(a)** coronal sections showing the location of the anterodorsal (AD), anteroventral (AV) and anteromedial (AM) subregions of the anterior thalamic nuclei (ATN) and **(b)** the largest (grey) and smallest (black) ATN lesions. (**c**) The number of ATN neurons (NeuN-positive) was substantially lower in all 17 ATN-lesion rats compared to 9 of the intact rats examined, which included four of the rats in the Sham-Trace group and five of the rats in the Sham-No Trace group. The cell count was made from a 1:4 series throughout the ATN and is therefore an underestimate. = median value, which was 3388 in rats with ATN lesions and 15339 in rats with sham lesions; **E** = the ATN-lesion rat that was excluded because it did not meet the a-priori criteria of at least 50% damage bilaterally with at least 25% damage within either hemisphere. **(d)** 2.5x magnification photomicrographs of NeuN staining in a sham rat and an ATN-lesion with an average lesion. Compared to the sham rat, there was clear reduction of cells as well as collapse in the tissue although AD and dorsal AV showed some sparing. Black dashed line indicates outline of the ATN.

### Spatial working memory in the RAM

As expected, both groups of ATN-lesion rats showed severely impaired performance when spatial working memory was tested in the 8 arm RAM (Fig. 3). The initial similarity in performance across groups is because most rats ran for 10 minutes but made relatively few arm entries at the start of testing. The number of errors made by Sham and ATN-lesion rats diverged on day two of training after which both ATN-lesion groups showed no evidence of improvement compared to both Sham groups (Group, F_3,27_=39.66, p<0.001; Group x Day, F_27,243_=4.75, p<0.001). The two sham groups did not differ, but they both differed markedly (p<0.001) from each of the ATN-lesion groups, which did not differ. Rats in both ATN-lesion groups also made fewer correct arm choices before the first error, while rats in both Sham groups progressively entered more arms prior to making an error as testing continued (Group, F_3,27_=27.53, p<0.001; each Sham group versus each ATN group, p<0.001; Group x Day, F_27,243_=2.21, p<0.001). Impaired spatial working memory can be regarded as a benchmark measure of ATN lesion effects (Aggleton and Nelson, 2015). As the two ATN-lesion groups showed a similar level of impairment in spatial working memory, subsequent comparisons on the non-spatial tasks are unlikely to be influenced by lesion differences between these two groups.

**Figure 3.**
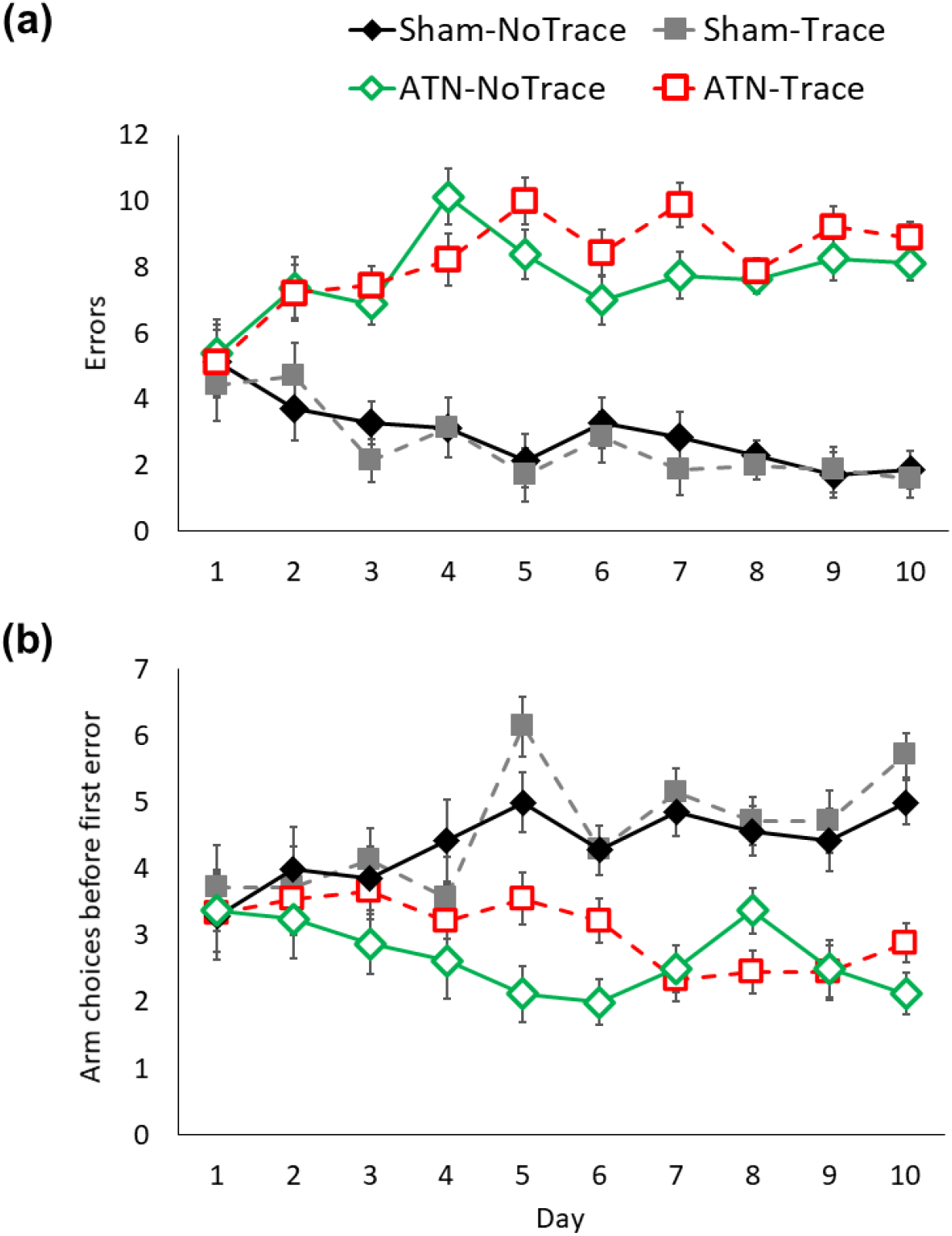
ATN lesions impaired spatial working memory in the radial arm maze. **(a)** Number of errors made per day and **(b)** number of choices before the first error. Mean +/-standard error. Sham-No Trace N=7; Sham-Trace N=7; ATN-No Trace N=8; ATN-Trace N=9.

### Non-spatial simple discrimination tasks

Acquisition of both the simple odour, and simple object, go-no go discrimination tasks is shown in Figure 4a and 4b. All rats rapidly acquired these tasks. The latency difference scores for the four groups did not differ in the simple odour discrimination task (Group, F_3,27_=0.97, p=0.42; Group x Day, F_15,135_=1.4, p=0.15). The groups took an average of 4-5 days to reach criterion in the simple odour discrimination task (Group, F_3,27_=1.99, p=0.13).

**Figure 4.**
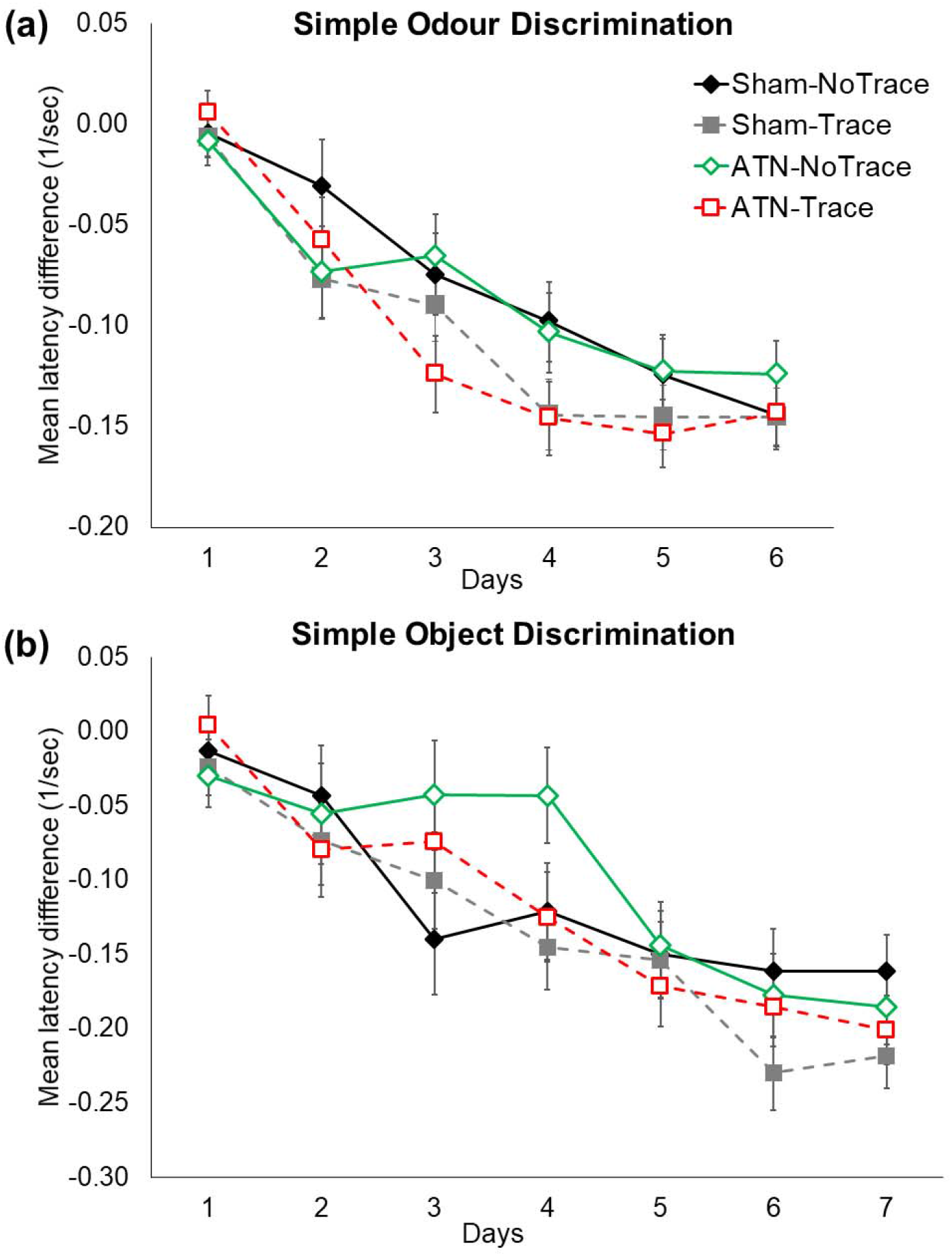
ATN lesions did not impair acquisition of either **(a)** the simple odour discrimination task or **(b)** the simple object discrimination task, which required an additional training day on average. The order of tasks was counterbalanced within each group. Mean latency is expressed as relative difference scores between non-rewarded and rewarded trials. The reciprocal latency (1/sec) was used because of the non-homogeneity of variance produced by raw latencies. Mean +/-standard errors. Sham-No Trace N=7; Sham-Trace N=7; ATN-No Trace N=8; ATN-Trace N=9.

The four groups also learned the simple object discrimination task without any significant differences (Group, F_3,27_=1.07, p=0.37; Group x Day, F_18,162_=0.97, p=0.49). Rats took an average of 5-6 days to reach criterion on the simple object discrimination task (Group, F_3,27_=0.30, p=0.82).

### Non-spatial paired associate tasks

#### Acquisition

The mean running latency during acquisition by the four groups is shown in Figure 5. Latencies during acquisition were carried forward for rats that reached criterion. The main findings were clear. The two Sham-lesion groups acquired their respective tasks, but not a single ATN-lesion rat showed task acquisition irrespective of the inclusion of a 10-second trace between the odour and object stimuli. The primary evidence for acquisition is whether the rats learned to inhibit their response to the object on the non-rewarded trials (Fig. 5a). On this measure, the two Sham groups progressively delayed responding to the object on non-rewarded odour-object pairings during training (i.e. showed reduced reciprocal latency scores). By contrast, all rats in both ATN-lesion groups instead showed increasingly faster response latencies as training progressed. That is, on non-rewarded trials, the ATN-lesion rats learned only to search more quickly under the object despite the odour-object pairing being incorrect. Note, the two ATN-lesion groups were, if anything, slower to respond than the Sham-lesion groups at the start of training on the non-rewarded trials, so general hyperactivity was not evident after ATN lesions in this task (Block 1 latency range: ATN-lesion 2.5-5.0 seconds, Sham-lesion 1.9-4.5 seconds; Block 10: ATN-lesion 1.4-3.0 seconds, Sham-lesion 5.5-7.6 seconds). Anova of latencies on the non-rewarded trials produced a significant main effect across the four groups (F_3,27_=22.63, p<0.001) and a significant Group x Block interaction (F_27,243_=26.27, p<0.001). The two Sham groups did not differ on this measure, and the two ATN groups did not differ, but both Sham groups differed from each of the ATN-lesion groups (p<0.001).

**Figure 5.**
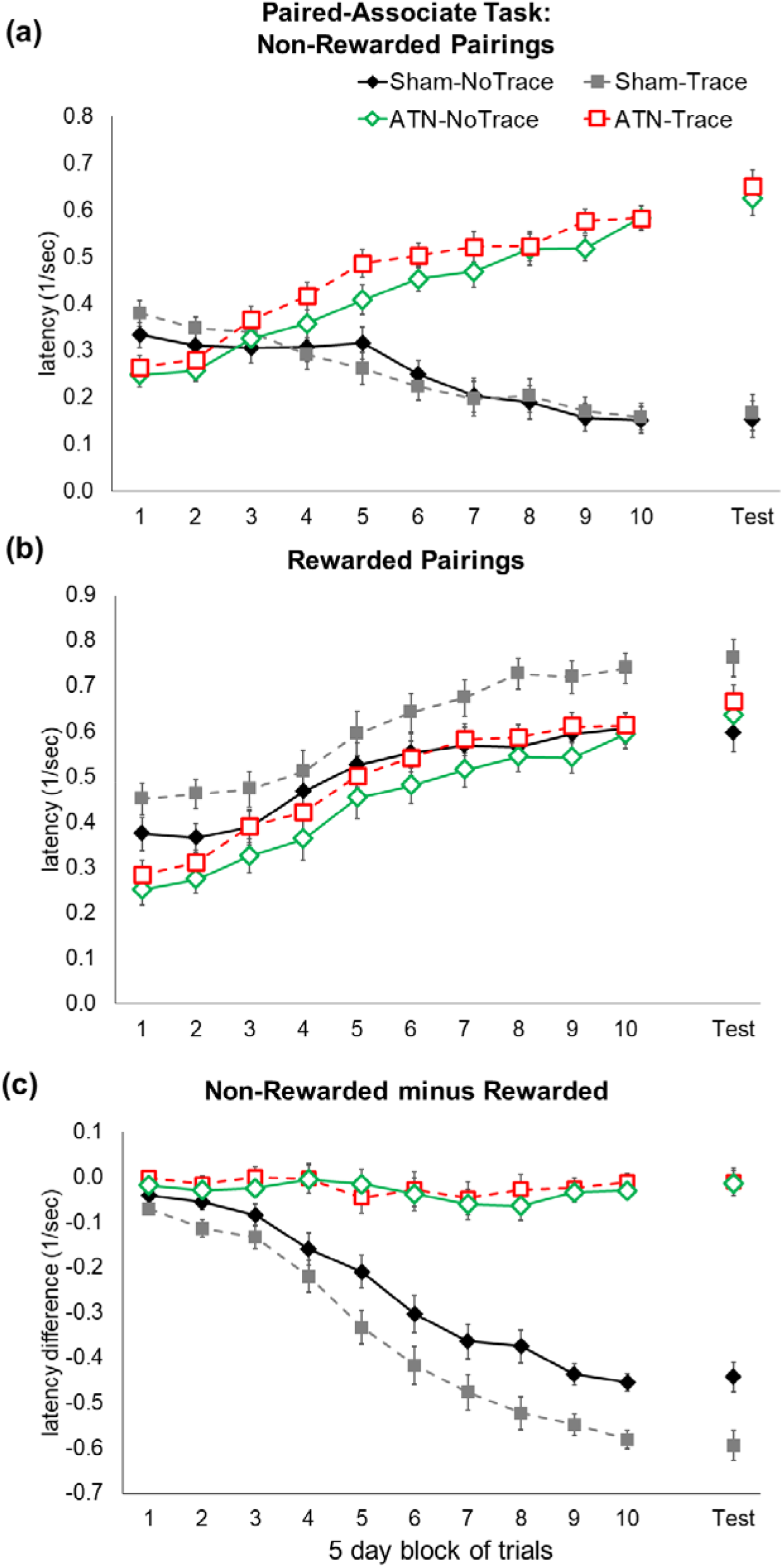
ATN lesions prevented acquisition of the odour-object paired association memory task, irrespective of the inclusion of a 10-second interval (trace) between the odour and object. Mean latency for (**a**) non-rewarded pairings and (**b**) rewarded pairings on the odour-object (no trace) and odour-trace-object (trace) paired-associate learning tasks. (**c**) Task acquisition and retention expressed as relative latency difference between non-rewarded and rewarded pairings for the four groups. The retention session was conducted 5 days after the rat either reached criterion or the end of the acquisition period. The reciprocal latency (1/sec) was used because of the non-homogeneity of variance produced by raw latencies. Mean +/- standard errors. Sham-No Trace N=7; Sham-Trace N=7; ATN-No Trace N=8; ATN-Trace N=9.

When the odour-object pairing was rewarded, all four groups showed increasingly faster response latencies (i.e. showed increased reciprocal scores) across blocks of trials (Block main effect, F_9,27_=75.78, p<0.001; Fig. 5b), irrespective of Group (Group x Block, F_27,243_=0.63, p=0.92; Fig. 5b). The Group main effect was again significant (F_3,27_=5.67, p=0.003), which in this instance was due to faster running speed by rats in the Sham-Trace group compared to both ATN groups (p<0.008) and the Sham-No-Trace rats (p=0.02). However, the remaining three groups showed similar response latency on the rewarded trials. Again, note that the two ATN-lesion groups showed slower responding at the start of training on the rewarded trials, compared to the Sham-lesion groups.

The direct comparison across latencies for non-rewarded and rewarded trials is shown in Figure 5c, expressed as latency difference scores. Figure 5c clearly shows that no learning occurred in either ATN-lesion group, whereas both Sham-lesion groups acquired the task (Group x Block interaction, F_27,243_=23.99, p<0.001). The latency difference score suggests that the Sham-Trace group acquired the task more strongly than the Sham-No Trace group, and the mean differences between these two groups were significant for the last three blocks of trials (p<0.01). This difference between the two sham groups, however, was primarily driven by their differences on trials with rewarded stimulus pairings (compare Fig. 5b with 5a).

#### Retention Test

All groups showed responding on the non-rewarded session at the 5-day retention test that was similar to their corresponding performance on Block 10 of training (Group x Block [Retention Test versus Block 10], F_3,27_=2.00, p=0.13; Fig. 5a). Despite the nonsignificant Group by Block interaction, there was an overall difference in latency across Block 10 versus the retention test (Retention versus Block 10, F_1,27_=7.22, p=0.01) that seems mostly due to ATN-lesion rats responding faster in the retention test. A significant group main effect (F_3,27_=73.13, p<0.001) was driven by Sham groups continuing to inhibit responding on the non-rewarded trials, unlike the ATN-lesion groups (each Sham group versus each ATN-lesion group, p<0.001).

For rewarded trials, all four groups showed similar mean response latencies in the retention test compared to latency in Block 10 of training (Block, F_1,27_=3.75, p=0.06; Group x Block, F_3,27_=1.02, p=0.39). Faster running speeds in the Sham-Trace group than all other groups was evident across Block 10 and the retention test (Group, F_3,27_=3.85, p=0.02; Sham-Trace versus all other groups, p<0.03).

The direct comparison of rewarded and non-rewarded trials on the retention test, expressed as a latency difference score (Fig. 5c), confirmed that all four groups maintained similar performance on this measure compared to their Block 10 acquisition performance (Group, F_3,27_=189.43, p<0.001; Block, F_1,27_=0.08, p=0.76; Group x Block interaction, F_3,27_=0.22, p=0.88).

#### Zif268 Expression

Our primary interest was to determine group differences in the regions of interest. Given the behavioural outcomes obtained, we do not report associations between Zif268 expression and performance as they would produce spurious correlations that are driven by the clear non-lesion versus sham status differences and small within-group variation on the primary measure of interest, that is, latencies on non-rewarded trials. Similarly, any associations evident in regions such as the retrosplenial cortex would again create an artificial correlation due to the marked group-level differences in Zif268 expression.

#### Prefrontal regions

Zif268 expression across the four groups in the prelimbic prefrontal cortex (A32V) and the anterior cingulate cortex (A32D; A24b; A24a) is shown in Figure 6. There was a significant Group main effect for Area 32V (F_2,37_=5.49, p<0.01; Fig. 6b). Here, the two ATN groups showed similar overall expression to each other (p=0.32), as well as the Sham-No Trace group (p>0.1), but both lesion groups showed lower expression than in the Sham-Trace group (p<0.02). The lower Zif268 expression in the Sham-No Trace group compared to the Sham-Trace group did not reach significance (p=0.06). There was also a significant main effect across the four Layers (F_3,81_=147.8, p<0.001), but no Group x Layer interaction (F_9,181_=1.15, p>0.3).

**Figure 6.**
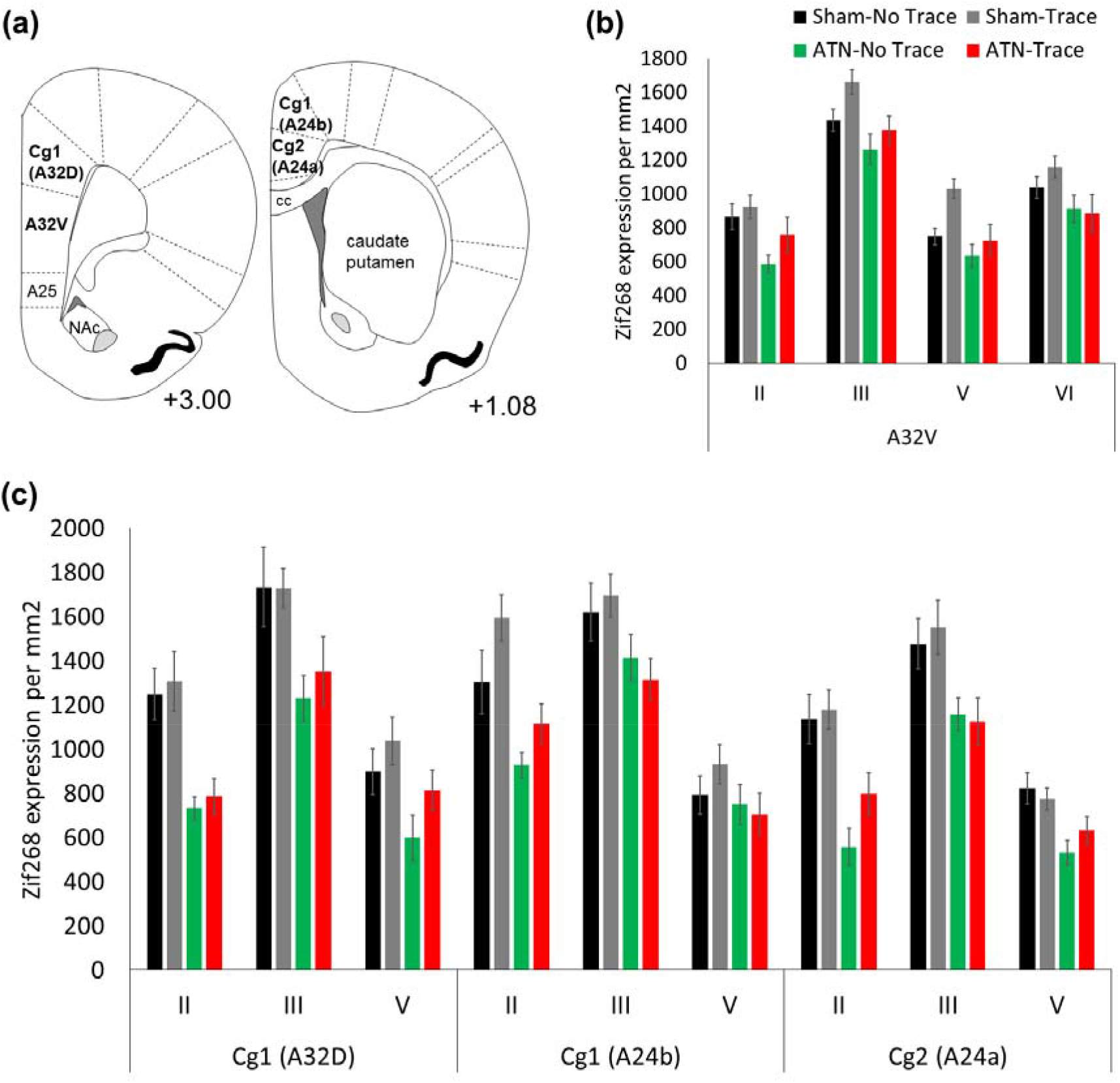
Zif268 expression assessed in the prefrontal cortex regions in the four groups after the 5-day retention test. The ATN-lesion groups showed decreased expression across the prefrontal regions, especially the superficial layers II and III. **(a)** Schematic diagram of +3.00mm and +1.08 from Bregma defining the prefrontal regions assessed. **(b)** Expression per mm2 in the prelimbic cortex (A32V) and **(c)** cingulate Cg1 (A32D and A42b) and Cg2 (A24a) regions (mean ± S.E). Sham-No Trace N=7; Sham-Trace N=7; ATN-No Trace N=8; ATN-Trace N=9.

For the anterior cingulate (Cg) regions (Fig. 6c), the two ATN-lesion groups showed lower Zif268 expression than both of the two Sham-lesion groups (Group F_3,27_=12.47, p<0.001; both Sham groups differed from each of the ATN groups, p<0.004, but not from each other, p=0.31). A significant Group x Layer interaction (F_6,54_=4.02, p<0.002) reflected larger differences between ATN and Sham groups for layer II and layer III (p<0.001) than for layer V, where the groups did not differ significantly (p>0.1). Expression differed across the three cingulate regions (Region main effect, F_2,54_=11.41, p<001), being lowest for A24a (posterior) compared to both A32D and A24b (p<0.002), and was highest for Layer III (Layer, F_2,54_=242.38, p<0.001) but the size of these differences varied between layers across the three cingulate regions (Region x Layer, F_4,108_=5.29, p<0.001). However, the Group x Region and Group x Region x Layer interactions were non-significant (all F<1.0).

#### Hippocampal and parahippocampal regions

Figure 7 shows the Zif268 expression for the four groups in hippocampal and parahippocampal regions. For the hippocampus, the dorsal CA1 and CA3 subregions, especially CA1, showed higher expression than the ventral hippocampal CA1 and CA3 (dorsal versus ventral, F_1,24_=541.4, p<0.001; CA1 versus CA3, F_1,24_=759.3, p<0.001; interaction between these two factors, F_1,24_=428.4, p<0.001). The most interesting finding, however, was that there was higher expression in the dorsal CA1 in the Sham-Trace group than in each of the other three groups (p<0.02), supported by a significant triple interaction for Group x [dorsal region versus ventral region] x [CA1 versus CA3], F_3,24_=6.09, p<0.003). For the other three groups, dorsal CA1 expression was the lowest in the ATN-Trace group (ATN-Trace versus Sham-No Trace and ATN-No Trace, p<0.03), while the two No Trace groups did not differ significantly (p=0.84). Analysis of the dorsal dentate gyrus (DG; only dorsal was examined) showed higher expression in the hilus than the granular cell layer (F_1,27_=386.1, p<.001), which was greater for the Sham-Trace and ATN-No Trace groups than for the Sham-No Trace and ATN-Trace conditions (Group x DG Subregion, F_3,27_=4.07, p<0.01). The ventral subiculum showed higher expression than the dorsal subiculum (F_1,27_=9.02, p<0.005), but there was no Group effect (F_3,27_=1.41, p=0.25) or Group x [dorsal versus ventral regions] interaction (F_3,27_=0.15, p=0.92). In the parahippocampal regions, the perirhinal cortex showed lower expression than the two entorhinal cortex areas (F_2,54_=16.2, p<0.001), but the groups did not differ at these sites (Group, F_3,27_<1.0; Group x Region, F_6,54_=1.8, p>0.1).

**Figure 7.**
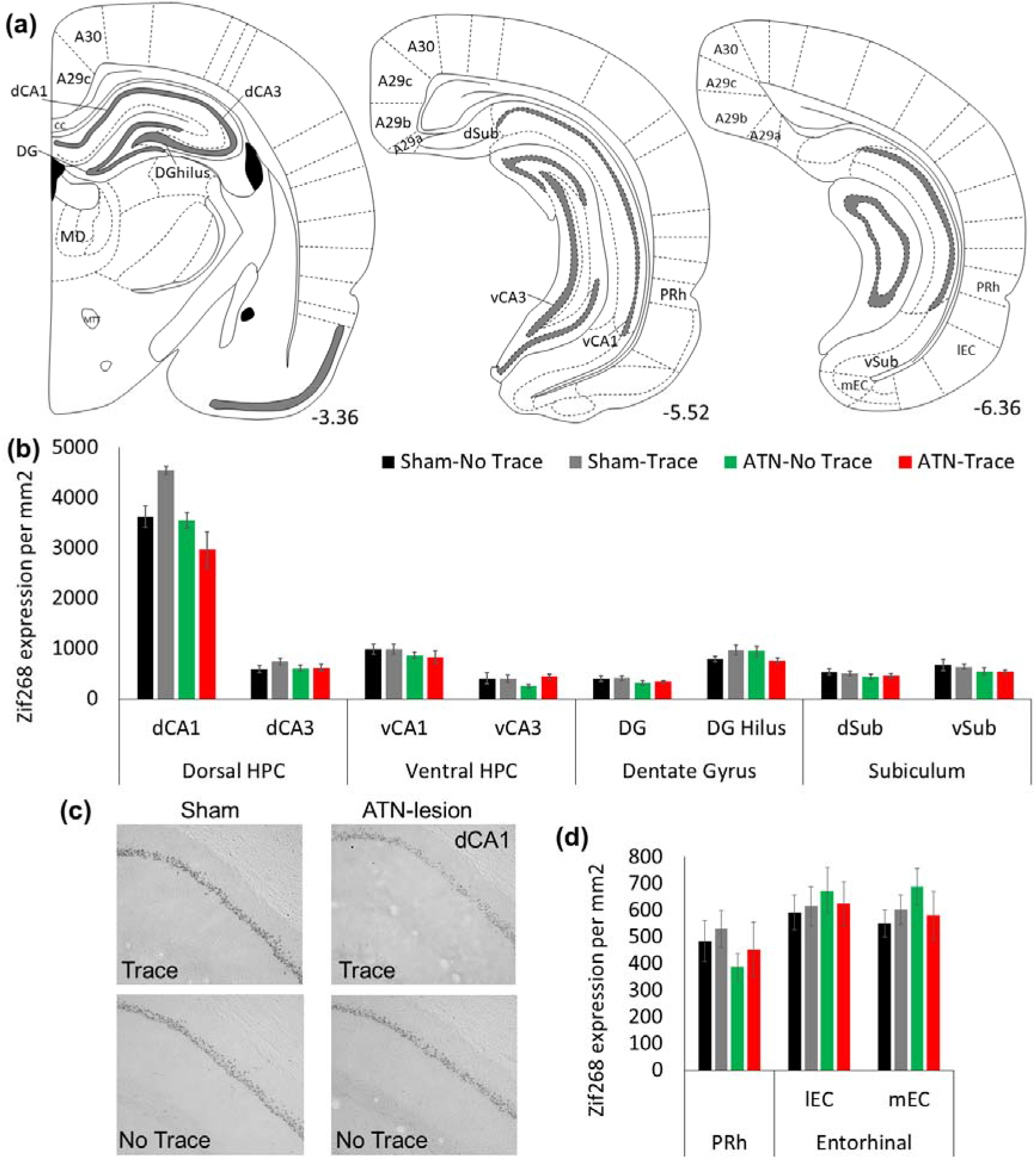
Zif268 expression assessed in regions of the hippocampal, subicular and parahippocampal regions in the four groups after the 5-day retention test. The Sham-Trace group showed higher Zif268 expression in the dorsal CA1. **(a)** Schematic figure of the regions analysed (numbers represent mm from Bregma). **(b)** Expression/mm2 in the dorsal and ventral hippocampal CA1 and CA3), dorsal dentate gyrus (granule cell layer and hilus), and dorsal and ventral subiculum. **(d)** Parahippocampal regions (perirhinal and lateral and medial entorhinal cortices). Mean ± S.E. (**c**) 10x magnification examples of Zif268 expression in the dorsal CA1 in the four groups (at ∼ −3.36mm from Bregma); examples show the rat with the median Zif268 value in each group. Sham-No Trace N=7; Sham-Trace N=7; ATN-No Trace N=8; ATN-Trace N=9.

#### Retrosplenial cortex

Figure 8 shows the Zif268 expression in the superficial and deep layers of Rga, Rgb and Rdg. The two ATN groups showed markedly lower expression across these three regions than was shown by the two Sham groups (Group main effect, F_3.27_=40.9, p<0.001). For the aggregated values across the three regions, the two Sham groups had higher mean Zif268 values than both ATN-lesion groups (p=0.0001) but the Sham-Trace group also showed higher levels than the Sham-No Trace group (p=0.04). The difference between Sham groups and ATN groups was smallest for the Rga region (Group x Region, F_6,54_=2.72, p<0.02). The level of Zif268 expression between groups also varied across layers (Group x Layer, F_3,17_=40.0, p<0.001). This interaction reflected differences between both Sham groups compared to both ATN groups that were larger in the superficial layers than in the deep layers. Nonetheless, the Sham versus ATN group effects were still significant in the deep layers (p<0.001). In addition, the Sham-Trace group showed higher expression in the superficial layers compared to the Sham-No Trace group (p=0.03), while expression in the deep layers did not differ between these two groups (p=0.23). There was no Group x Region x Layer interaction (F_6,54_=1.0, p=0.43).

**Figure 8.**
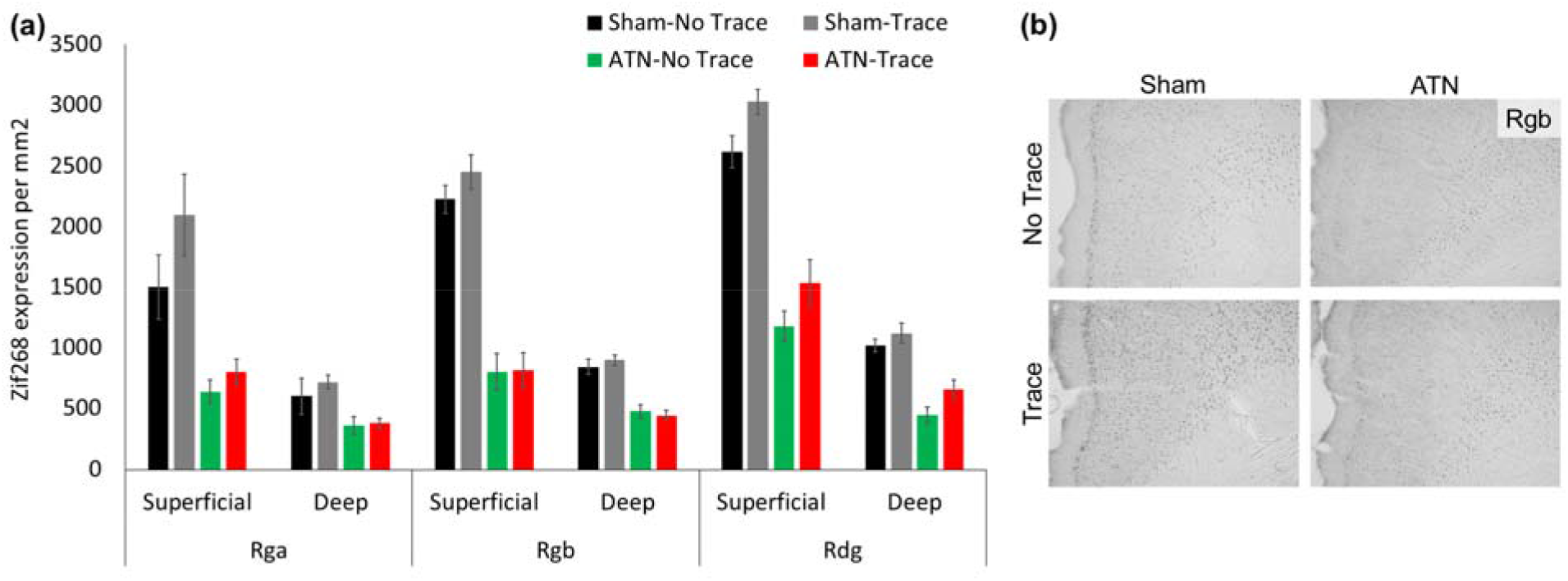
Zif268 expression in the retrosplenial cortex for the four groups after the 5-day retention test. ATN lesions produced in a significant reduction of Zif268 expression across all retrosplenial regions. (**a**) Expression per mm^2^ in the granular (Rga and Rgb) and dysgranular (Rdg) regions of the retrosplenial cortex (mean ± S.E). (**b**) 10x magnification examples of Zif268 expression in the four groups showing superficial and deep layers of the Rgb at ∼ - 3.36mm from Bregma (the median rat in each group). For schematic representation of retrosplenial regions see Figure 7. Sham-No Trace N=7; Sham-Trace N=7; ATN-No Trace N=8; ATN-Trace N=9.

#### Auditory control cortex

No differences were found in the auditory (control) cortex (Group, F_3,27_=1.2, p=0.35).

## Discussion

The aim of this study was to examine the effects of ATN lesions on non-spatial paired-associate memory and the influence of an explicit delay (i.e. a 10 second trace) between the presented odour and object stimuli. Evidence of impaired paired-associate memory after ATN lesions has previously been reported only when one of the paired components required the processing of distal spatial cues (Dumont et al., 2014; Gibb et al., 2006; Sziklas & Petrides, 1999). We had anticipated that the effects of ATN lesions on non-spatial paired-associate memory in our task would be most evident when an explicit trace procedure was used. This was because it has been suggested that CA1 lesions only impair non-spatial paired-associate memory when a 10-second “trace” is used (Kesner at al., 2005) and the microstructural integrity of CA1 neurons is reduced by both ATN lesions (Harland et al., 2014) and by mammillothalamic tract lesions that cause ATN dysfunction (Dillingham et al., 2019). Moreover, the key examples of non-spatial memory impairment after ATN lesions examined temporal discriminations among multiple object or odour items presented within a single block of trials (Wolff et al., 2006; Dumont & Aggleton, 2013). However, none of 17 rats with ATN lesions showed evidence of acquisition of the odour-object paired-associate task, including those not trained with an explicit delay between the non-spatial stimuli. Despite an extended period of training, the ATN-lesion rats were unable to show inhibited responding to the non-rewarded odour-object pairings. General hyperactivity in the ATN-lesion rats does not seem a feature of this impairment, because they showed slower responding than the sham-lesion rats during the initial stages of acquisition. The Sham-Trace group showed shorter latencies than the other three groups on the rewarded trials, but this difference may reflect increased anticipation of reward when restrained for a 10-second delay rather than a measure of faster acquisition by this group. Faster acquisition would be expected to be reflected by response inhibition on the non-rewarded trials but the two sham-lesion groups did not differ on this measure.

The failure to learn the paired-associate memory tasks after ATN lesions does not appear to be due to poor inhibition or impaired sensory processing. The ATN-lesion rats demonstrated rapid acquisition in both the simple object discrimination task and the simple odour discrimination task, which was equal to that shown by sham-lesion rats. The task demands for these simple discriminations were identical to the paired-associate task and used the same apparatus. For the same reason, the paired-associate deficit after ATN lesions is also unlikely to be due to a simple failure of attention to the individual stimuli used. ATN lesions impair the ability to learn an attentional-set and facilitate extradimensional shifts, but do not change sustained attention or behavioural flexibility (Chudasama & Muir, 2001; Wright et al., 2015; Kinnavane et al., 2019). The rapid acquisition of the simple discriminations in the current runway task contrasted with slower acquisition when we trained a previous group of rats on an open circular platform to learn a simple odour discrimination and, especially, a simple object discrimination (Bell, 2007). So, it is possible that the use of a runway and the explicit reduction of distracting spatial cues, plus the active interaction with the object to search for food, facilitated attention to the non-spatial stimuli in the current study.

The severity of the lesion impairment for learning the association between odour and object stimuli suggests that this task is strongly dependent on the integrity of the ATN. This evidence is at odds with the suggestion that paired-associate impairments after ATN lesions require the use of multi-modal spatial stimuli (Dumont et al., 2014; Nelson, 2021). One possible explanation for the difference between the outcome in the current study and that of Dumont et al. (2014) is that discrete non-spatial stimuli in biconditional discrimination learning tasks, such as specific objects or odours, may pose a greater attentional demand on the establishment of a unique integrated representation compared to the use of a general local context, such as the thermal, visual or tactile local environment. In a similar way, the profound susceptibility of spatial paired-associate tasks to impairment after ATN lesions may also rely on the integration of relational spatial cues in combination with a salient discrete cue, because acquisition of a simple spatial discrimination per se was only partially impaired (Dumont et al., 2014; Gibb et al., 2006). One unexpected finding is that rats with ATN lesions showed no deficit when they were required to select a particular location in a cross-maze on the basis of a conditional visual cue at a choice-point (Sziklas & Petrides, 2007). In that situation, however, the conditional relationship was determined by a single salient cue presented a single place that had no ambiguity or embedded association with the different positions of the correct spatial location. This contrasts with a complete failure by ATN-lesion rats when they had to learn an object-place association in which they had to select one of two correct objects on the basis of their associated location (Sziklas & Petrides, 1999).

The finding that ATN lesions produced a profound deficit in odour-object paired-associate memory, irrespective of the presence of a 10-second trace between the stimuli, adds to our earlier evidence of a deficit of paired-associate memory when the object and odour were presented simultaneously on a cheeseboard platform (Bell, 2007). Together, this suggests that the memory deficits after ATN lesions on odour-object paired-associate memory is an additional example that ATN lesions do not always mirror the pattern of conditional associative memory deficits produced by lesions to the hippocampal system (Sziklas & Petrides, 2004, 2007). While ATN lesions can cause greater spatial memory impairments than fornix lesions (Warburton & Aggleton, 1999), or deficits in object-place and geometric discrimination tasks that are not found with fornix lesions (Aggleton, Poirier, Aggleton, Vann, & Pearce, 2009; Sziklas, Lebel, & Petrides, 1998), there is less evidence that ATN lesions can produce severe memory impairments in tasks that are generally unaffected by lesions of the hippocampal formation. With respect to non-spatial arbitrary associations, the evidence that the hippocampus is only critical when a 10-second trace is used between the two stimuli was derived by comparing acquisition in two different tasks. Gilbert and Kesner (2002) reported that large hippocampal lesions did not impair object-odour paired-associate memory when tested on a cheeseboard platform in which the two stimuli were presented simultaneously. In a similar runway to ours, but with an object presented before exposure to odourised sand that might contain a reward, Kesner et al. (2005) showed that dorsal CA1 lesions but not CA3 lesions produced a deficit when using a 10-second trace condition. Neither we nor Kesner and colleagues examined the effects of hippocampal lesions in the runway task without the trace condition. So, we cannot be sure that rats with hippocampal lesions would be unimpaired under the no-trace condition when trained in the runway using our procedures. Nonetheless, our study extends the predicted association between CA1 function and temporal processing in a non-spatial paired-associate task by finding increased Zif268 expression in dorsal CA1 in the Sham-lesion group trained using a 10-second trace relative to the Sham-lesion No Trace group. By contrast, the mean Zif268 expression in the dorsal CA1 was lowest in the ATN-lesion Trace group. There was also evidence, albeit weaker, that the trace condition in sham-lesion rats was associated with increased Zif268 expression in the superficial layers of the retrosplenial cortex. The pattern of different behavioural performance in the lesion and non-lesion groups at retention made it inappropriate to examine the association between variations in performance and variations in Zif268 expression. As in previous studies (Aggleton & Nelson, 2015; Perry et al., 2018) the strongest effect was a marked reduction in IEG expression after ATN lesions in the retrosplenial cortex, especially the superficial layers. It seems likely that this finding is due to the loss of or diminished activity in the direct inputs from the ATN to the RSC (Barnett et al., 2021).

There has been growing awareness that ATN lesions may exert influences beyond spatial memory impairments (Nelson, 2021; Wolff et al., 2006). One example is when ATN lesions slows acquisition of non-spatial attentional-set learning, which may be due to a functional relationship between the ATN and the mid-cingulate regions of the cingulate cortex rather than medial prefrontal cortex connections (Wright, et al., 2015; Bubb et al., 2021). To our knowledge, however, this attentional-set task has not been examined with hippocampal lesions in rats. A clearer example of a dissociation between ATN lesions and lesions of the hippocampus is that only the former injury impairs attentional processes associated with latent inhibition (Nelson et al., 2018). It is possible that an impoverished ability to establish the relevance or predictiveness of stimulus-stimulus associations provides a unifying account not only of attentional-set learning and latent inhibition (see Nelson et al., 2018) but also instances of impaired paired-associate learning after ATN lesions. Rather than ascribing the role of the ATN from the perspective of either hippocampal (for space and time) or frontal (for attentional) processes, the broader implication is that the ATN may support memory processing by actively orchestrating attention to certain classes of stimulus-stimulus associations and their representation across multiple brain structures (Leszcynski & Staudigl, 2016). Precisely how to characterise the classes of impaired memory remains an experimental challenge for the future. What is clear, from the current study, is that explanations based only on a spatial / non-spatial dichotomy are unable to account for profound memory deficits that can be found in both domains after ATN lesions.

Our findings bring the effects of ATN lesions in rats closer into line with paired-associate memory impairments in clinical amnesia after injury to the mammillary body – ATN axis (Rempel-Clower, Zola, Squire, & Amaral, 1996; Squire et al., 2020). There is, however, a clear difference in the slow acquisition of paired associate memory in intact rats and the rapid acquisition of paired-associate memory in humans with intact memory systems. Paired-associate tasks are assumed to reflect episodic-like memory by measuring the ability to form unique representations of multiple stimuli rather than memory for individual components (Eichenbaum & Fortin, 2009; Crystal & Smith, 2014). In our study, however, the intact rats required 4 to 5 weeks and over 300 training trials before there was clear evidence of acquisition, which suggests that the task could be more rule-based or semantic-like in these rats. This limitation could be circumvented in future work by first training intact rats on one or more non-spatial paired-associate tasks before testing the acquisition of a new odour-object pairing after ATN lesions. In this way, the general rule of making an association would already be established and, perhaps, the rate of acquisition would then be relatively rapid for a new task in intact rats. In addition, temporary chemogenetic or optogenetic manipulations of the ATN could provide an opportunity to investigate the impact that these nuclei have on retarding acquisition rather than preventing acquisition or on their impact on retention rather than acquisition. It would also be informative to learn whether many or only some neural projections from the ATN support this example of non-spatial paired-associate learning. Neuroanatomical evidence suggests a different pattern of neural connections with limbic and cortical memory structures among the three ATN nuclei, that is, the anterodorsal nuclei, anteroventral nuclei and the anteromedial nuclei (Bubb et al., 2017; Lomi et al., 2021; Nelson, 2021). In addition, these ATN component nuclei have different molecular and electrophysiological characteristics that may underpin different behavioural functions (Jankowski et al., 2013; Roy et al., 2021; Roy et al., 2022; Safari et al., 2020). Lesions or genetic manipulations of the individual nuclei of the ATN could provide insight as to whether paired-associate memory relies on one or more of the AD, AV or AM. It may be the case that the impact of ATN lesions on this task preferentially involves frontal brain regions and / or the retrosplenial cortex and their respective involvement in rule-based and knowledge-based systems rather than event-based memory (Hunsaker & Kesner, 2018). These issues could also be addressed using contralateral disconnection lesions involving the ATN, as this experimental approach has been used to successfully demonstrate the system-wide influence of the ATN-hippocampal axis in spatial tasks (Dumont et al., 2010; Warburton et al., 2000, 2001).

The present study provides unequivocal evidence that ATN lesions produce substantial impairments in non-spatial paired-associate learning and memory, irrespective of the presence of an explicit temporal component. Extensive training demonstrated no evidence of learning in these ATN-lesion rats. The fact that there was relatively little to minimal damage to the immediately adjacent intralaminar or mediodorsal thalamic regions suggests that these impairments were specific to the ATN lesion. Evidence from these non-spatial paired-associate tasks suggests a new perspective on the role of the ATN as a critical node within the ‘hippocampal-diencephalic-cingulate’ memory network (Bubb et al., 2017). This strengthens the view that the ATN do not operate primarily as a relay for hippocampal information (Wolff & Vann, 2019). Instead, the ATN may actively control the generation of some classes of arbitrary memory representations in the brain.

## Acknowledgements

This research was supported by University of Canterbury equipment and research grants and Early Career support (JJH) from Brain Research New Zealand – Rangahau Roro Aotearoa.

## Author Contributions

JJH and JCDA: funding, concept and design, writing and editing drafts, statistical analysis and interpretation of data; JJH: conducting experiments.

## Data accessibility

Will be available on Mendeley

## Abbreviations

A29a: area 29a of the granular retrosplenial cortex
A29b: area 29b of the granular retrosplenial cortex
A29c: area 29c of the granular retrosplenial cortex
A30: area 30 of the dysgranular retrosplenial cortex
A32D: area 32D of the anterior cingulate cortex
A32V: area 32V of the medial prefrontal cortex
A24a: area 24a of the anterior cingulate cortex
A24b: area 24b of the anterior cingulate cortex
ATN: anterior thalamic nuclei
AD: anterodorsal thalamic nuclei
AM: anteromedial thalamic nuclei
AV: anteroventral thalamic nuclei
CA1: cornu ammonis 1
CA3: cornu ammonis 3
cc: corpus callosum
Cg1/Cg2: cingulate area 1 / 2
DG: dentate gyrus
DG Hilus: hilus of the dentate gyrus
dSub: dorsal subiculum
IEG: immediate early gene
lEC: lateral entorhinal cortex
MD: mediodorsal thalamic nuclei
mEC: medial entorhinal cortex
MTT: mammillothalamic tract
NAc: nucleus accumbens
PRh: perirhinal cortex
Re/Rh: reuniens/rhomboid nuclei
Rga: dysgranular a retrosplenial cortex
Rgb: dysgranular b retrosplenial cortex
Rdg: granular retrosplenial cortex
V: ventricle
vCA1: dorsal cornu ammonis area 1
vCA3: dorsal cornu ammonis area 3
vSub: ventral subiculum.

